# OpenGRF: Predicting Ground Reaction Forces and Moments During Daily Living Activities in OpenSim

**DOI:** 10.1101/2025.09.27.678739

**Authors:** Andrea Di Pietro, Francesca Di Puccio, Luca Modenese

## Abstract

**Background and Objectives:** Ground reaction forces and moments (GRF&Ms), typically measured using force plates, are key inputs for musculoskeletal simulations. OpenSim currently lacks a tool to predict GRF&Ms directly from kinematics. This study presents OpenGRF, an OpenSim-based tool designed to estimate GRF&Ms and the centre of pressure (CoP) from joint kinematic data and validates its performance against force plate recordings.

**Methods:** The proposed methodology integrates calibrated foot–ground contact probes with an optimization framework based on computed muscle control, while CoP is computed accounting for both kinematic and dynamic contributions. For validation, a scaled FullBodyModel (37-DoF without muscles) was created for seven healthy adults performing six trials each of level walking, stair ascent, and stair descent, for a total of 126 marker-based trials. GRF&Ms predictions were compared to reference force plate data using normalized RMSE (nRMSE), Pearson correlation coefficients (ρ), CoP error, and Statistical Parametric Mapping (SPM)

**Results:** Results showed high accuracy for vertical GRF (nRMSE ≤1.5%, ρ ≥0.94), particularly during level walking, and good accuracy for anterior–posterior GRF (nRMSE 4.4–6.1%, ρ = 0.81–0.91). Medio–lateral GRF was less reliable, especially in stair tasks (nRMSE up to 11.2%, ρ down to 0.48). Free moments were the most challenging quantity to predict across all tasks (nRMSE up to 28%). In contrast, ankle moments were predicted with high fidelity (nRMSE ≈1.7%, ρ ≈0.98). Median CoP errors were 21–23 mm, with largest discrepancies during double support.

**Conclusions:** OpenGRF enables physics-consistent estimation of GRF&Ms and CoP directly from kinematics, achieving the highest accuracy for vertical GRFs and predicting ankle moments that closely match those obtained from force plate measurements.

## 1. Introduction

The prediction of ground reaction forces (GRFs) and moments (GRMs, collectively GRF&Ms) in the absence of force plates (or other experimental measurement) is of great interest to the biomechanical field [1–8]. It enables dynamic analysis, such as estimating joint torques for biomechanical and sports analyses and, in the musculoskeletal (MSK) modelling domain, could enable the use of muscle-driven simulations from kinematic data. Various methods have been used to estimate GRF&Ms from kinematic data, ranging from applying machine learning techniques (e.g.[9–17]) to solving the dynamic Equations of motion in foot-ground models through optimization methods [1,6]. Among machine learning approaches, Oh et al. [13] adopted an artificial neural network to solve the statically indeterminate problem of the double support phase during gait while, more recently, Cordero-Sànchez et al. [17] took advantage of a similar approach to predict GRFs from running kinematics. Alternatively, Shahabpoor et al. [18] exploited a nonlinear system identification method (NARMAX) to estimate GRFs during walking from body segments accelerations obtained via inertial measurement units (IMUs). Meanwhile, Liu et al. [19] trained two bi-lateral long short-term memory neural networks to predict 3D GRFs during stair negotiations using whole-body marker-based kinematics as the input. However, machine learning-based methods, require substantial amounts of high-quality training data to achieve and improve accuracy, which are generally available for gait but not necessarily for other activities of daily living.

On the other hand, regarding optimization-based methods, Fluit et al. [1] proposed a methodology to estimate the GRF&Ms that addresses the indeterminacy during double-support by incorporating them into the static optimization algorithm that solves muscle recruitment. Skaals et al. [20] further validated this approach on dynamic motor tasks, enhancing the foot-ground contact formulation, which then was adapted by Konrath et al. [21] for estimating knee adduction moment using inertial motion capture, while Kloeckner et al. [22] utilised it to predict GRF&Ms in children with cerebral palsy during gait. The optimization method proposed by Fluit et al. [1], however, is implemented in the commercial MSK modelling software Anybody [23], and it is not currently available in other programs.

Currently, the open-source software OpenSim [24] does not provide an embedded resource for GRF&Ms prediction. The two OpenSim-based tools, OpenCap [25] and Moco [26], among their various capabilities, offer potential approaches to estimate these quantities, but so far, to authors’ best knowledge, their application has been demonstrated only for a limited set of motor tasks, with accuracy and generalizability still under investigation. Consequently, achieving accurate and generalizable prediction of GRF&Ms within OpenSim remains an open challenge for both marker-based and markerless applications (e.g. OpenSense [27] and OpenCap [25]) .

Building on this, we propose a novel methodology, called OpenGRF (freely available at https://simtk.org/projects/opengrf), to estimate GRF&Ms from kinematic data by leveraging the optimization tools within the OpenSim platform [24]. We validated this approach using a marker-based dataset of daily living activities (level walking, stairs ascent and descent). For these activities, the centre of pressure (CoP) and GRF&Ms were predicted and compared against corresponding measured ground-truth data. To further strengthen the validation, joint kinetics derived from predicted versus measured GRF&Ms were also compared. Validation was performed using quantitative metrics and a statistical parametric mapping analysis (SPM) [28].

## 2. Materials and methods

### 2.1 Experimental data

Motion capture data were collected from seven healthy young individuals (age: 27.1 ± 2.1 years, mass: 61± 13.2 kg, height: 167.3 ± 11.3 cm) at the “Biodynamics Lab” of Imperial College London in the context of a previous musculoskeletal modelling project approved by the institution’s ethics committee. Each participant performed six repetitions of level walking, stair ascent, and stair descent activities, yielding a total of 126 trials.

A full-body marker set was used, with marker placement following the International Society of Biomechanics (ISB) recommendations for both the lower and upper extremities [29,30]. The marker trajectories were captured using a 10-camera Vicon system (Vicon Motion Systems, Oxford, UK) at 100 Hz. Ground reaction forces and moments (including CoP) were synchronously recorded with three Kistler force plates (Kistler Instrumente AG, Winterthur, Switzerland) at 1000 Hz.

### 2.2 Input data preparation

For each participant, the FullBodyModel of Rajagopal et al. [31] having 37 degrees of freedom (DoFs) was used. The left and right metatarsophalangeal joints were locked at the reference position reducing the model to 35 DoFs. The model was scaled to each participant’s anthropometrics using the Automated Scaling Tool [32] on a static trial. This procedure yielded a mean RMSE tracking error of 4 ± 0.1 mm between experimental and model markers, across subjects. The scaled model was subsequently used to run the Inverse Kinematic tool and compute joint angles for all activities. The resulting kinematic data were low-pass filtered with a 6 Hz cut-off Frequency for further OpenSim analyses.

### 2.3 Algorithm description

The algorithm implemented in OpenGRF was developed using the MATLAB API of OpenSim 4.5. It requires as input a scaled MSK model (“.osim”) and a joint kinematics file (“.mot”), and generates a “.mot” file containing the GRF&Ms applied to each foot, along with the corresponding CoP locations (see Figure 1 and 2). The algorithm consists of three main phases: 1) Contact analysis of the foot-ground pose, 2) GRF&Ms estimation using a Computed Muscle Control (CMC)-based approach, and 3) CoP and free moment calculation during the single-stance phase based on the solved dynamics of the system (Figure 1 and 2). Simulations were run on a laptop equipped with an Intel i7 processor (@ 2.7 GHz) and 16 GB RAM requiring on average 5 minutes to simulate each second of input kinematics.

**Figure 1.**
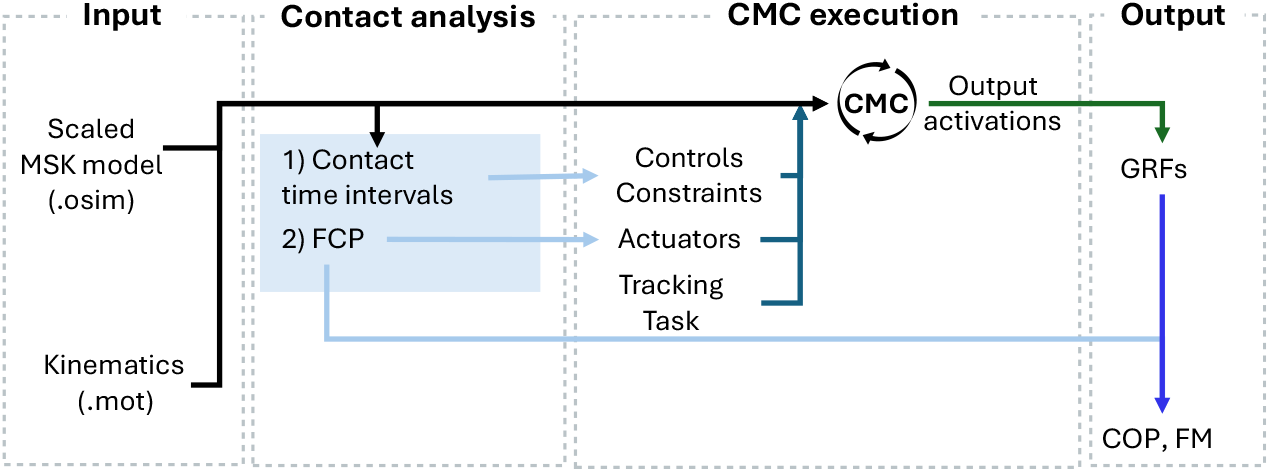
Relationship between the input (scaled MSK model and kinematics files) and output (ground reaction forces (GRFs), free moment (FM) and centre of pressure (CoP)) values in OpenGRF.

**Figure 2.**
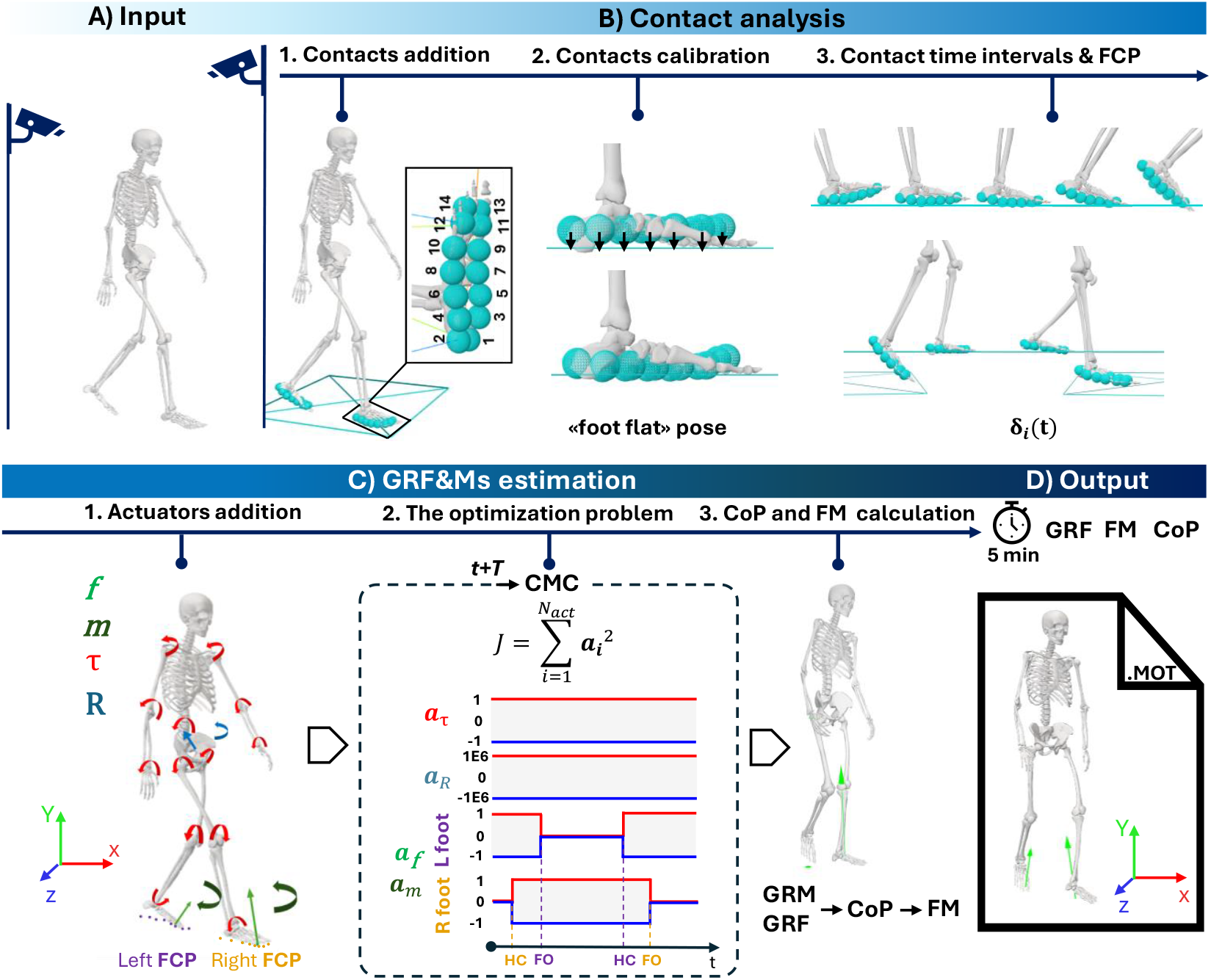
OpenGRF workflow: A) MSK model and kinematics used as input; B) Contact analysis: B1) Addition of the 14 contact probes on the models feet and contact planes; B2. calibration of contact probes B3) Analysis of the task computing contact time intervals and Foot Contact Point (FCP); C) GRF&Ms estimation: C1) Addition of ideal actuators (coordinate, external and residual) to the model; C2) Execution of the cyclic Computed-Muscle-Control (CMC) tool with constrained actuators activations C3) calculation of the Centre of Pressure (CoP) and free moment (FM) during single stance based on the obtained dynamics; D) predicted GRFs, FM and CoP stored in “.mot” file as output.

#### 2.3.1 Foot-ground contact analysis

The first phase of the proposed methodology identifies contact between the model foot segments and the ground, which requires a two-step preparatory procedure (Figure 2, 1^st^ row): 1) 14 spherical contact probes are added to the model foot segments as contact probes, and 2) a calibration phase refines their locations with respect to the ground using the available limb kinematics. This analysis determines the intervals of foot-ground contact, including the timing of single- and double-stance phases. It also provides the Foot Contact Points (FCPs), offering an initial estimate of the CoP location.

##### Contact probes definition and calibration procedure

Before conducting any analysis, a contact plane is added for each foot using the OpenSim ground frame as the default reference. A user-defined plane with height *y*_*0*_ is employed for the stair negotiation tasks. This plane is activated only when both the toe and the calcaneus centres of mass (CoM) lie above its height and their absolute speeds are sufficiently low (typically the 35% of their maximum values over the entire movement), ensuring reliable contact detection.

Two rows of spherical contact probes (radius: 2 cm) are added to each foot, one medially and the other one laterally, spanning from the posterior calcaneus to the middle phalanges (Figure 2, 1^st^ row). Probe positions are scaled with the model in the medio-lateral and anterior-posterior directions, while their centres are initially placed on an arbitrary horizontal plane. Their placement along the cranial-distal direction affects the extent of foot-ground penetration; therefore, calibration is required to account for the variable offsets arising from footwear or barefoot conditions.

The calibration procedure consists of the following steps (Figure 2, 1^st^ row):

1. For each foot, identify the time frame corresponding to the minimum vertical location of the calcaneus CoM and vertical acceleration [33] (“foot flat” instant).
2. Adjust the centres of the contact probes so that they uniformly intersect the contact plane at the identified foot-flat instant with a constant penetration of 7 mm, a default value set to avoid sudden and unrealistic contact loss.
3. Determine the maximum foot-ground penetration across the simulated task.
4. Adjust each contact probe position along the cranial-distal direction by the difference between the maximum and the adjusted foot-flat penetration, ensuring that the maximum depth of all spheres remains consistent across the trial.

Contact probes are only employed to compute contact time intervals, distinguishing also between single- and double-stance phases, and FCPs as described below.

##### Estimation of Contact Time Intervals and Foot Contact Point

The contact time intervals are identified as the time periods when at least one sphere has a non-zero penetration (*δ*_i_) with the contact plane. After calibration, *δ*_i_ is computed as the depth of each sphere relative to contact plane (Figure 2, 1^st^ row).

During contact, the FCP for each foot is computed as the weighted centroid of the probe centres projected onto the contact plane, with the foot-ground penetrations as weights, i.e.:

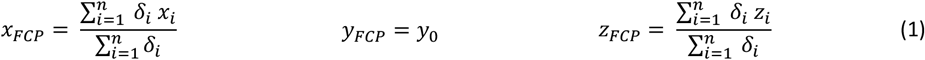

where *n*=14 is the number of spheres per foot, *x*_i_ and *z*_i_ are the coordinates of the centre of each contact sphere and *y*_0_ is the level of the contact plane (16 cm in this study). FCPs provide an approximation the CoPs during double support phases.

#### 2.3.2 GRF&Ms estimation

In musculoskeletal analyses, when the human body is treated as a whole system, Newton’s equations can be applied to estimate the resultant external force and moment, provided that the segmental kinematics and inertial properties are known. As detailed in the Appendix C, this formulation produces a single force applied at a specified point along with a single resultant moment. In single-support phases, these correspond to the ground reaction force and moment of the contact foot. A challenge arises during double support, where partitioning the resultant force and moment between the two feet becomes indeterminate. To address this issue, additional criteria or methodological approaches are required. In this study, we propose a modified version of the computed muscle control (CMC) tool available in OpenSim [34].

The CMC algorithm [34,35] generates dynamic simulations that track experimental kinematics using static optimization to determine muscle (or, more generally, actuator) recruitment. In its standard implementation, experimental GRFs are provided as inputs to the CMC tool In the proposed approach, muscles are disabled, while ideal actuators are incorporated between each calcaneus and the ground, to estimate unknown GRF&Ms. Specifically, three point-actuators, located at the previously identified FCP on each foot, generate the GRFs (one for each global axis), while three torque actuators generate the GRMs, resulting in six actuators per foot (12 in total). Each actuator produces an instantaneous force/torque equal to the product of its optimal magnitude with its level of activation (between -1 and 1) and are constrained to be active during the contact time intervals and inactive outside of those (see Appendix A and B)

Additionally, the model is actuated by a set of ideal actuators that provide generalised forces at the active model joints (29 coordinate actuators) and at the pelvis frame (6 residual actuators) to facilitate simulation convergence. In a simulation, the coordinate actuators generate any torque required to balance the joints, whereas the optimizer, minimizing the cost function (Eq. 3), tends to favour the engagement of the external actuators, which are characterized by very high optimal force resulting in consistently low activations, rather than the residual actuators, which are designed in the opposite way (see Appendix B).

##### The optimization problem

According to the standard CMC algorithm, in a first step a proportional-derivative feedback controller is applied to compute the desired generalized accelerations 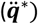 expressed by the following equation (Eq. 2) [34]:

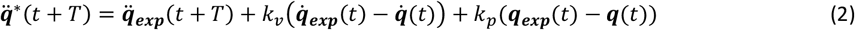

Where ***q*** and 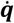 are, respectively, the vectors representing the generalized coordinates and velocities of the model; 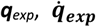 and 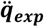 are respectively the experimental generalized coordinates, velocities and accelerations computed using inverse kinematics and considered the values to track. The default feedback gains *k*_*v*_ and *k*_*p*_, relating to velocity and positions errors respectively, critically damp numerical fluctuations. The look-ahead window (*T*) determines the time instant (*t* + *T*) at which 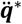 is calculated. In this study, *T* was set equal to the sampling rate of the marker and kinematic data (0.01 s).

In the second phase, the modified static optimization estimates the actuator activations, specifically:

- Coordinate actuators activations (***a***_***τ***_)
- Residual actuators activations at pelvis (***a***_***r***_)
- External forces (***a***_***f***_) and moments (***a***_***m***_) actuators activations.

In this case, the cost function of the optimization problem becomes (Eq. 3):

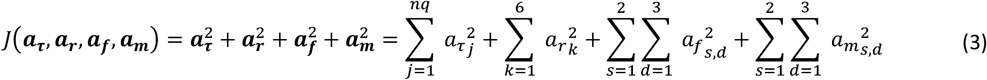

where the index *j* represents the model’s joint coordinates, excluding the pelvis DoFs indicated by *k*; *s* indicates the body side (left and right foot) while *d* refers to the GRF&Ms components.

The CMC ‘fast target’ tracking formulation ensures the difference between computed 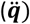 and desired generalized accelerations 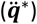 to be within a certain small interval (10^-5^ by default), effectively acting as additional constraints in the optimization problem [36]. In this process, the GRF&Ms are estimated because the optimization design variables ***a***_***f***_ and ***a***_***m***_ influence 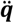 through the external loads they generate, as expressed in Eq. 4:

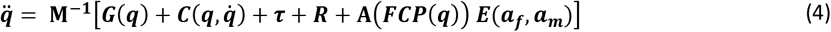

where **M** is the model mass matrix, ***G*** the vector of gravitational forces, ***C*** the vector of the centripetal and Coriolis forces, ***τ*** the vector of coordinate actuator forces, ***R*** the vector of residual forces, **A** the matrix expressing FCP locations as a function of the generalized coordinates (***q***), and **E** the vector of GRF&Ms. As noted earlier, ***a***_***f***_ and ***a***_***m***_ are subject to the time-varying constraints and are switched on or off depending on whether the ground contact is detected (Figure 2, 2^nd^ row).

As third step, the Forward Dynamics tool integrates the equations of motion up to *t* + *T*. The optimization and integration are then executed cyclically with time increments *T*. Finally, we applied a low-pass filter (Butterworth at 6 Hz in the current study) to the predicted GRF&Ms and exported them to an OpenSim-readable ‘.mot’ file (Figure 2, 2^nd^ row).

Due to the tracking nature of this methodology and the inclusion of residual actuators in the model, both the tracking errors and the magnitude of the pelvic residuals were quantified and assessed against thresholds recommended in previous literature [37].

#### 2.3.3 CoP and Free Moment calculation

The computation of the CoP location requires different considerations for the single- and double-support phases. As detailed in the Appendix C, during the single support, the system dynamics is fully determined, and the GRFs can be considered as applied at the CoP rather than the FCP. Therefore, CoP can be analytically derived from the GRFs applied at FCP and GRMs:

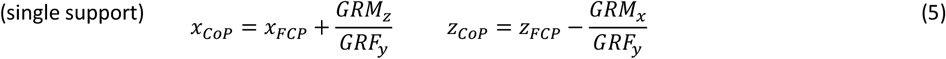

In contrast, during double support the system becomes statically indeterminate, and the FCPs are taken as approximations of the CoPs:

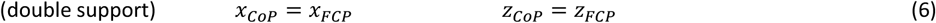

The transition between single and double support phases and vice versa, shifting from a determinate to an approximated solution and back, may introduce sudden gaps in the CoP location. To ensure continuity in the CoP trajectories across both phases, a weight is introduced in Eq.s 5, similarly to [1], leading to Eq.s 7 (see Appendix D):

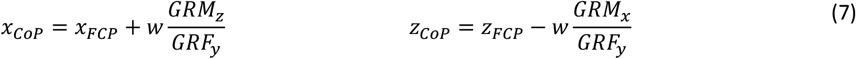

Where *w* is a quintic spline, ranging from 0 to 1, which enables a smooth transition over the first and last 20% of the single stance phase (Figure 3). Notably, the CoP vertical component (y direction) always lies on the contact plane (*y*_*CoP*_ = *y*_*FCP*_ = *y*_0_).

**Figure 3.**
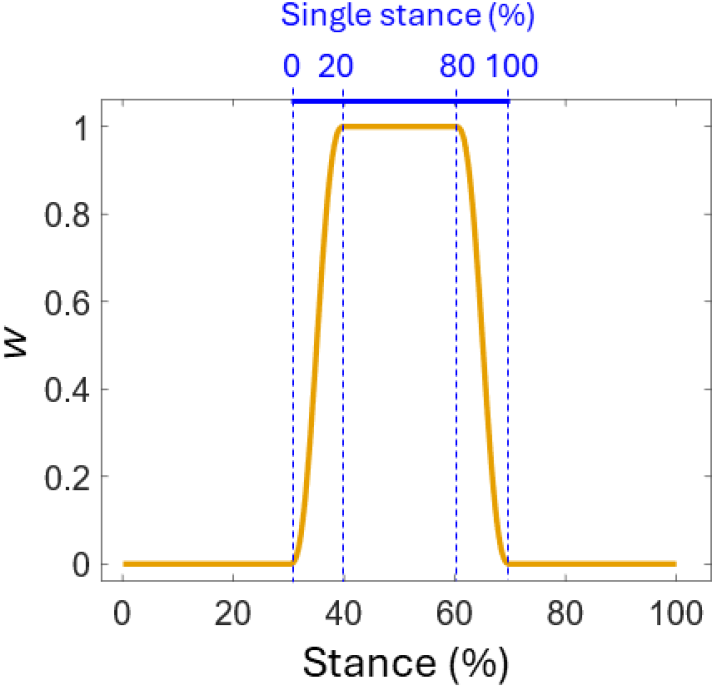
Representation of the weight function (w) for the dynamic-based CoP trajectories during single stance (%) that becomes null outside the single stance phase.

Accordingly, for single support phases (where w>0), the GRMs reduce to a single vertical component, commonly referred to as *free moment* (*FM*) (Eq. 8):

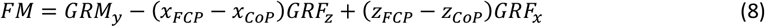

### 2.4 Predictions assessment and statistical analyses

For validation purposes, we assessed not only the GRF&Ms, but also their propagated effects on lower-limb joint kinetics at the ankle, knee and hip. Across all trials, predicted GRF&Ms were compared with their ground-truth data from the force plates using normalized Root Mean Squared Error (nRMSE) and Pearson’s correlation coefficient (ρ). The CoP error was quantified at each time frame as the distance (mm) between predicted and experimental positions.

Additionally, Statistical Parametric Mapping [28,38] (SPM) was performed using the *spm1d* toolbox in MATLAB R2024a (The MathWorks Inc. (2024), Natick, Massachusetts) to identify stance-phase intervals in which predicted and measured GRF&Ms were statistically different. After confirming the non-normal distribution of residuals, a two-tailed nonparametric 1D paired t-test was conducted at the 0.05 significance level. Bias, defined as the mean difference between prediction and ground-truth values for each component, was then calculated over the intervals identified as systematically different.

Joint kinetics obtained from measured and predicted GRF&Ms were compared using the same metrics.

## 3. Results

Using OpenGRF, CMC-based simulations for all participants and considered tasks were successfully performed and an estimation of the GRF&Ms obtained. The quality of the input kinematics was assessed through the IK errors (RMSE: 14±4, 23±3, 18±2 mm and maximum marker error: 38, 120 and 46 mm on a marker on the thigh obtained for level walking, stairs ascent and stairs descent, respectively), following the recommended thresholds for level walking [37]. The quality of the simulations was verified ensuring that the kinematic tracking errors were generally under 2 degrees, so compatible with the values suggested in the literature [37] (see Appendix E) and the residual actuators contribution to the simulation was also negligible (<5 N and <5 Nm, see Appendix F).

Among GRFs, the vertical (VE) GRF component was consistently the most accurate (Table 1). During level walking (Figure 4), it achieved the lowest error (nRMSE: 1.4 ± 0.4%) with correlations up to ρ = 0.99, and it remained highly accurate in stair ascent (nRMSE: 1.5 ± 0.6%, ρ: 0.94 ± 0.05) (Figure 5) and descent (nRMSE: 1.1 ± 0.6%, ρ: 0.96 ± 0.05) (Figure 6). The anterior-posterior (AP) component showed moderate accuracy (nRMSE: 3–8% and ρ: 0.81–0.96), whereas the medio-lateral (ML) component was less reliable in stair negotiation tasks (nRMSE up to 11.2 %, with ρ as low as 0.48). In contrast, the FM throughout the tasks showed reduced agreement, with nRMSE equal to 27 ± 9.4% in level walking and ρ as low as 0.4 ± 0.22 in stairs descent.

**Table 1.** Summary of Ground Reaction Forces (GRF) and Free moment (FM) and joint moments prediction assessment in terms of ρ (coefficient of correlation) and nRMSE (Normalized root mean square error) in terms of mean (standard deviation) per each motor task. AP=anterior-posterior, ML=medio-lateral, VE=vertical, FM= free moment, FE=flexion-extension, AA=ab-adduction, IE=internal-external.

| Task | Metric | GRF and Free Moment |  |  |  | Joint torques |  |  |  |  |
| --- | --- | --- | --- | --- | --- | --- | --- | --- | --- | --- |
|  |  | AP | VE | ML | FM | Ankle FE | Knee FE | Hip FE | Hip AA | Hip IE |
| Level walking | $\rho$ | 0.91<br>(0.07) | 0.94<br>(0.03) | 0.73<br>(0.14) | 0.63<br>(0.26) | 0.98<br>(0.01) | 0.72<br>(0.20) | 0.78 (0.07) | 0.87<br>(0.05) | 0.85<br>(0.10) |
|  | nRMSE | 4.6 (1.6) | 1.4 (0.4) | 5.9 (2.4) | 27.7<br>(9.4) | 1.7 (0.4) | 6.4 (1.9) | 7.1 (1.4) | 3.3 (0.6) | 5.3 (1.1) |
| Stairs Ascent | $\rho$ | 0.81<br>(0.11) | 0.94<br>(0.05) | 0.68<br>(0.2) | 0.47<br>(0.22) | 0.95<br>(0.03) | 0.95<br>(0.03) | 0.54 (0.21) | 0.83<br>(0.08) | 0.79<br>(0.11) |
|  | nRMSE | 6.1 (1.9) | 1.5 (0.6) | 7.8 (3.1) | 21.9<br>(14.1) | 2.6 (0.9) | 5 (2.1) | 11 (3.3) | 5.4 (1.6) | 10 (5.2) |
| Stairs Descent | $\rho$ | 0.90<br>(0.08) | 0.96<br>(0.03) | 0.69<br>(0.20) | 0.4<br>(0.22) | 0.89<br>(0.09) | 0.71<br>(0.24) | 0.61 (0.19) | 0.86<br>(0.13) | 0.83<br>(0.13) |
|  | nRMSE | 4.4 (1.9) | 1.1 (0.6) | 7.6 (3.6) | 23.2<br>(8.6) | 3 (2.1) | 8.3 (4.2) | 12.1 (5) | 4 (2.9) | 6.7 (3.1) |

**Figure 4.**
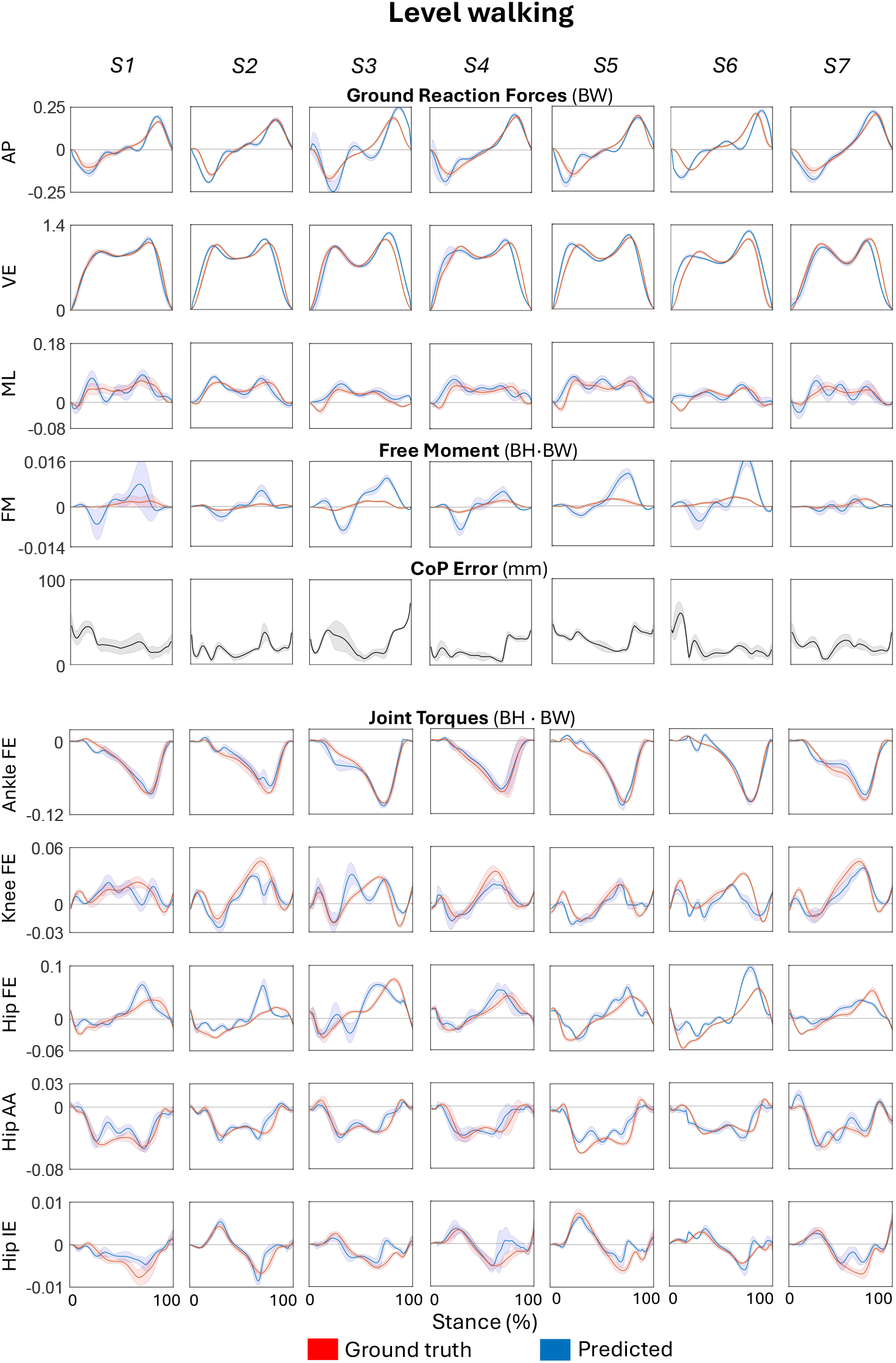
Results (mean ± SD) of Ground Reaction Forces (GRF) represented in Body Weight (BW), free moment (FM) and joint torques represented in body height · body weight (BH·BW) predicted (blue) against ground truth (red) for each of the 6 trials performed by the 7 subjects during level walking. Centre of pressure (CoP) errors shown in millimetres (mm). AP=anterior-posterior, ML=mediao-lateral, VE=vertical, FM= free moment, FE=flexion-extension, AA=ab-adduction, IE=internal-external.

**Figure 5.**
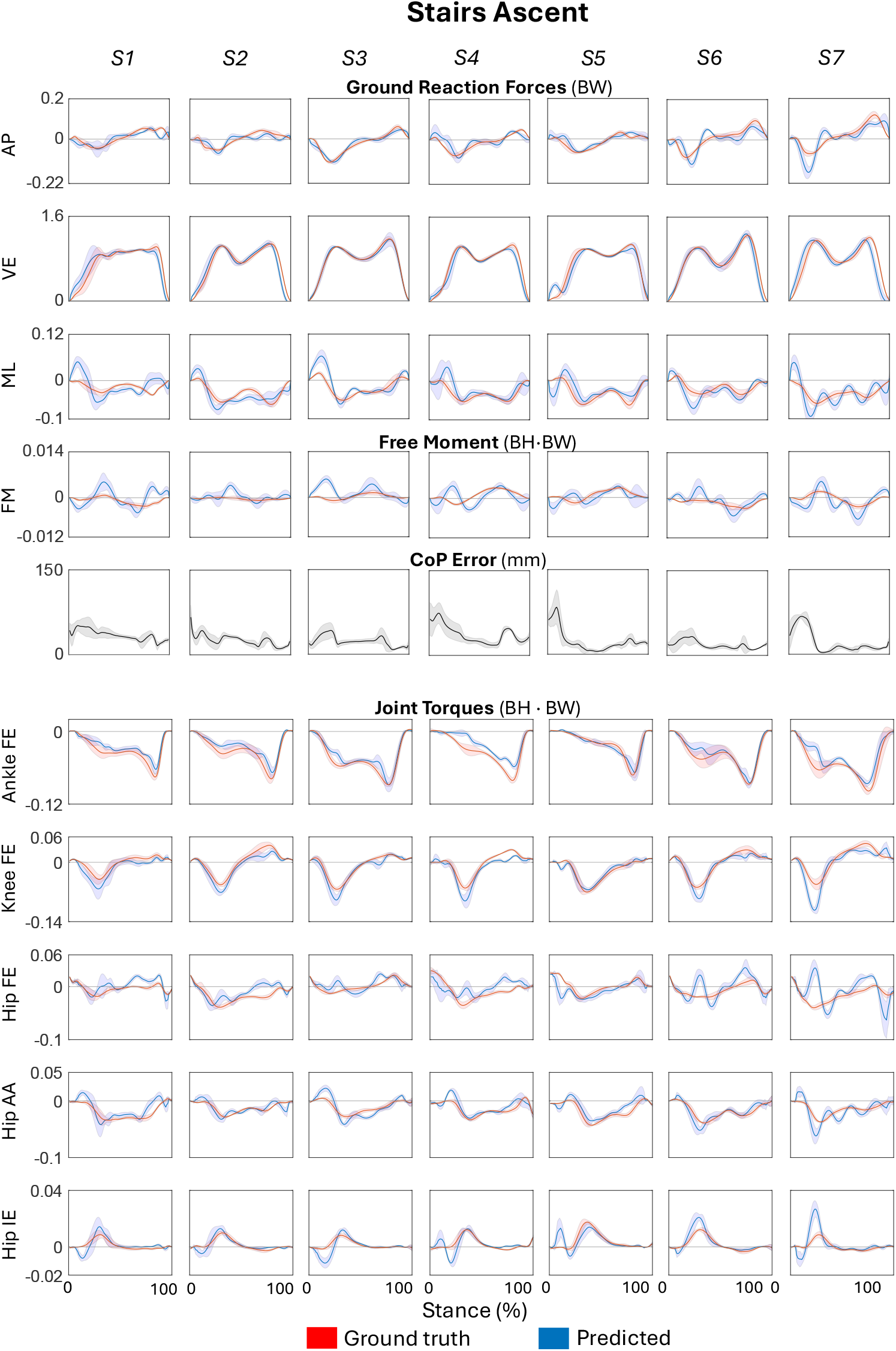
Results (mean ± SD) of Ground Reaction Forces (GRF) represented in Body Weight (BW), free moment (FM) and joint torques represented in body height · body weight (BH·BW) predicted (blue) against ground truth (red) for each of the 6 trials performed by the 7 subjects during stairs ascent. Centre of pressure (CoP) errors shown in millimetres (mm). AP=anterior-posterior, ML=medio-lateral, VE=vertical, FM= free moment, FE=flexion-extension, AA=ab-adduction, IE=internal-external.

**Figure 6.**
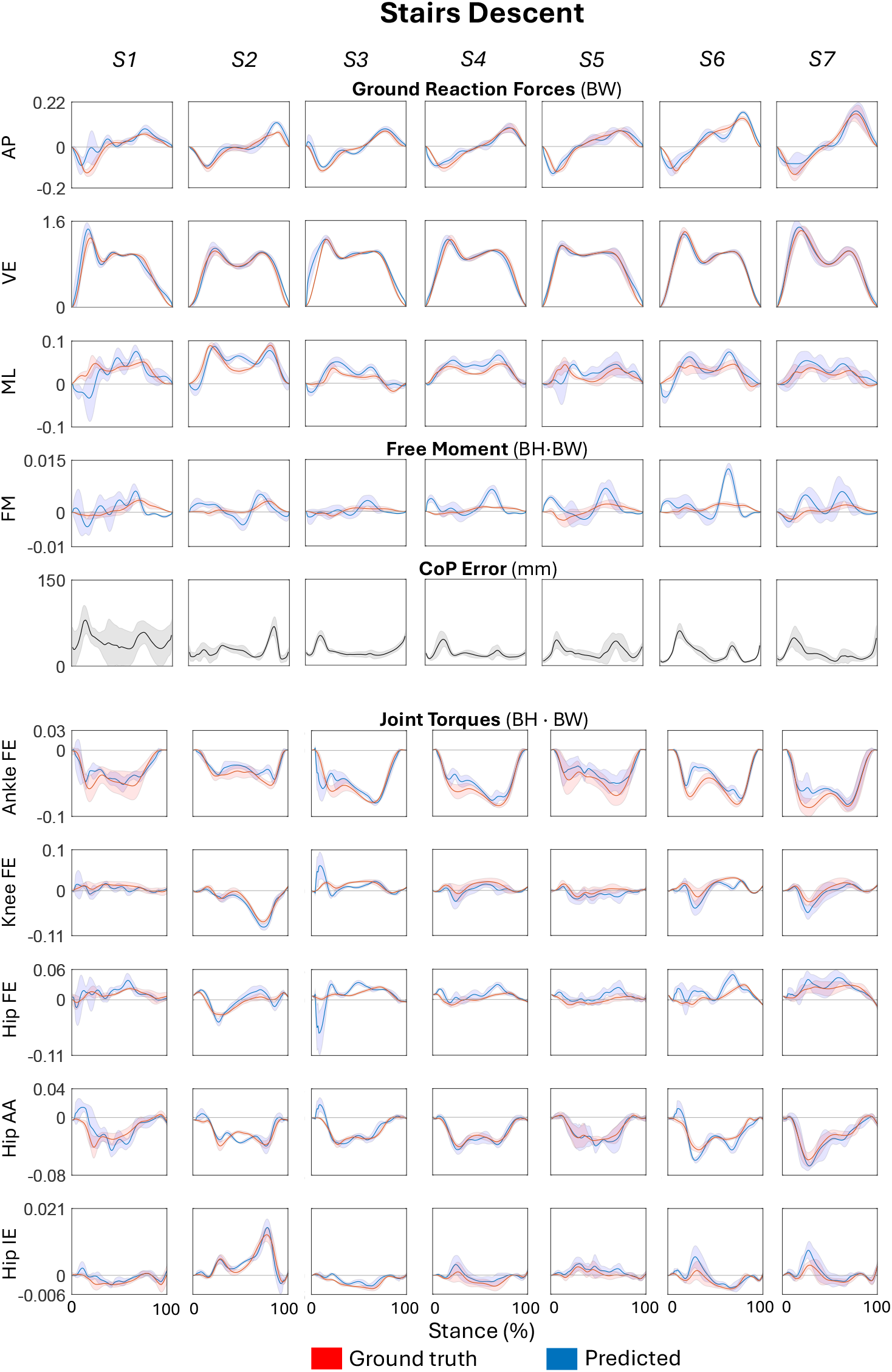
Results (mean ± SD) of Ground Reaction Forces (GRF) represented in Body Weight (BW), free moment (FM) and joint torques represented in body height · body weight (BH·BW) predicted (blue) against ground truth (red) for each of the 6 trials performed by the 7 subjects during stairs descent. Centre of pressure (CoP) errors shown in millimetres (mm). AP=anterior-posterior, ML=medio-lateral, VE=vertical, FM= free moment, FE=flexion-extension, AA=ab-adduction, IE=internal-external.

**Figure 7.**
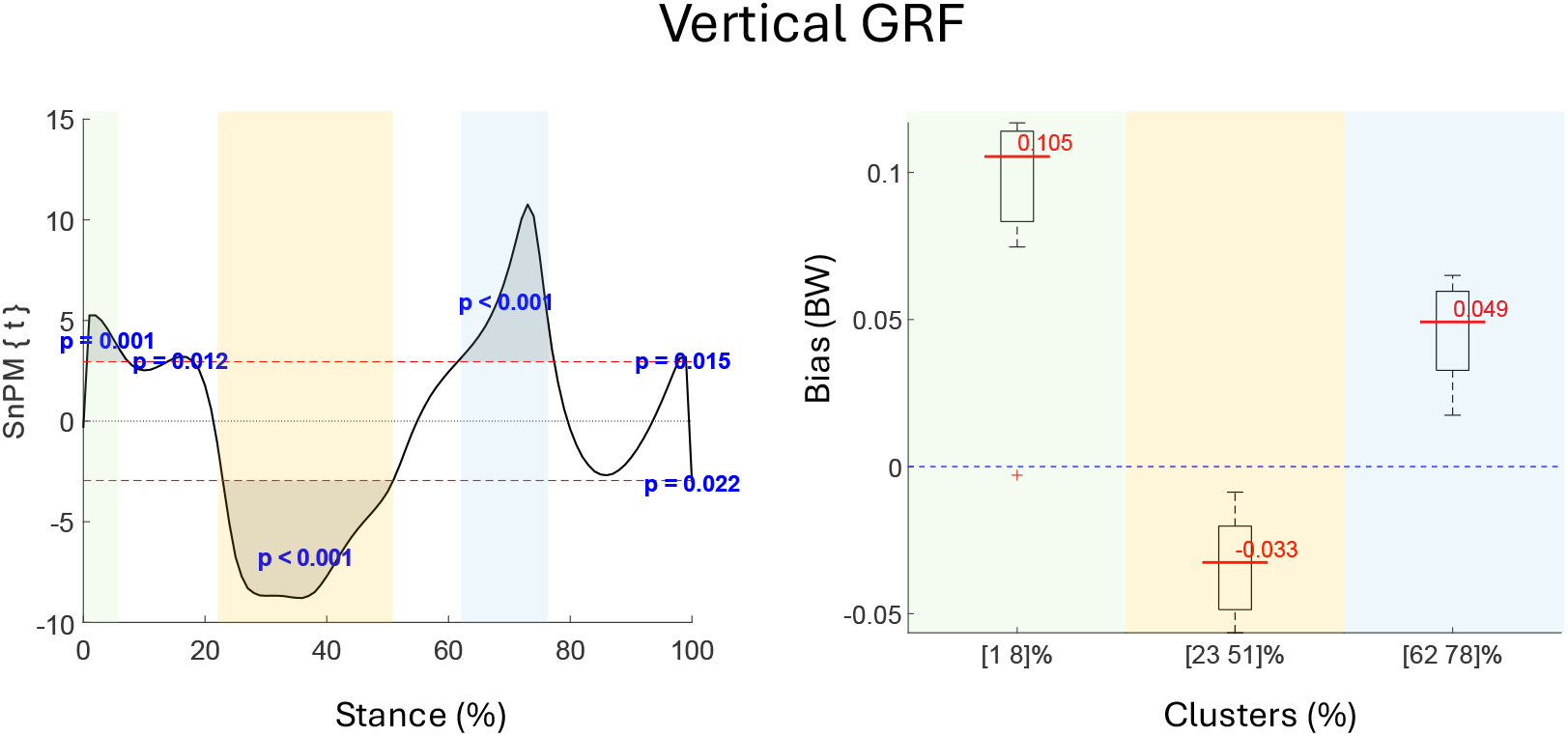
Example of the representation of result from SPM analysis: On the left there is the result of SPM analysis consisting of the two-tailed 1D paired t-test between the 42 (6 trials per 7 subjects) pairs (predicted vs ground truth) of curves; the red dashed lines represent the critical value associated with the 5% significance level beyond which the null hypothesis is rejected with the related p-value (in blue). On the right there are the boxplots per each of the clusters (each with a different color) detected as systematically different (with ρ<0.001) within the related interval of the stance phase. In each boxplot the continuous red line represents the median bias in that specific interval.

**Figure 8.**
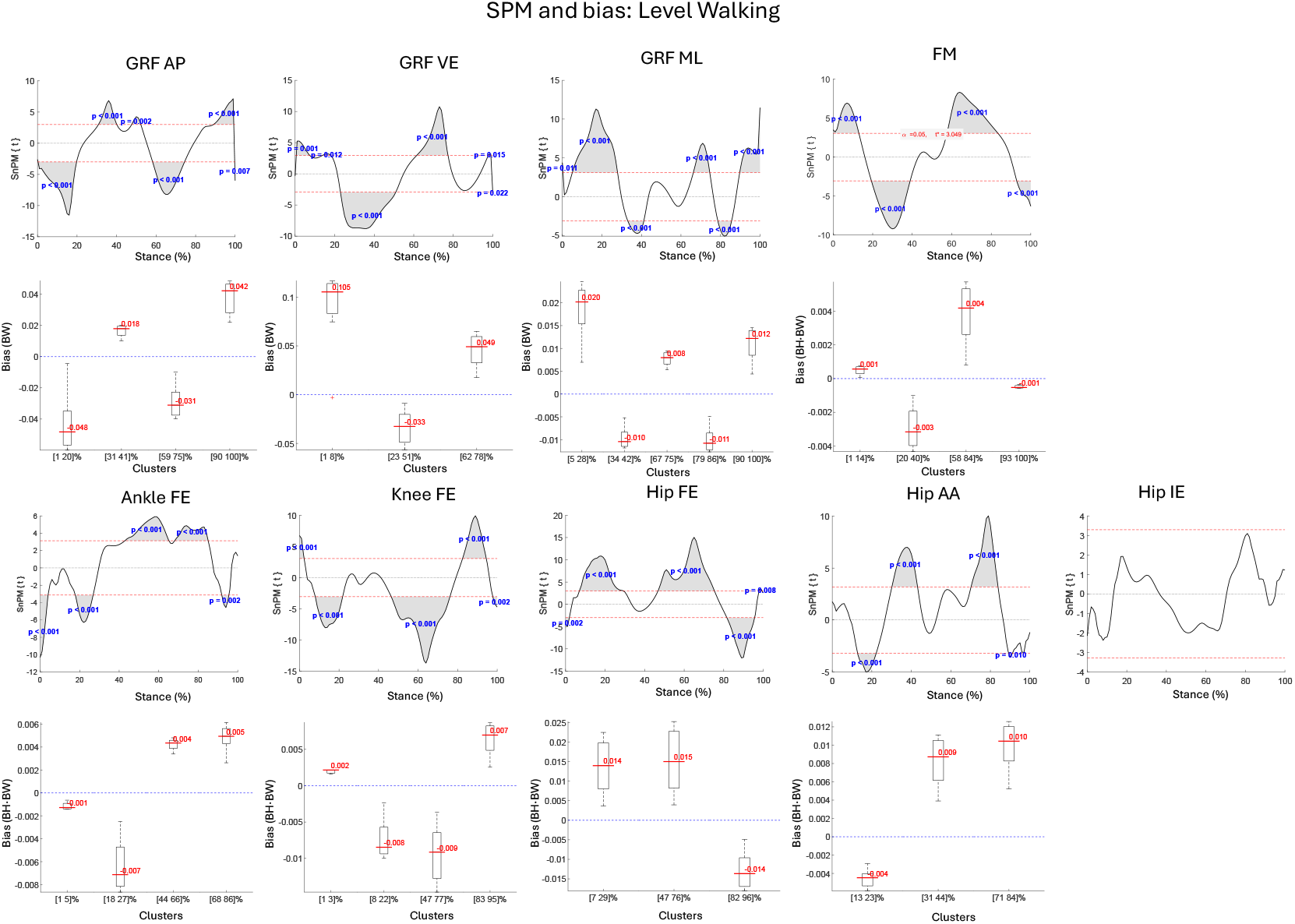
Results of SPM analysis between predictions and ground truth and related boxplots of biases for stance intervals with systematic differences during level walking.

**Figure 9.**
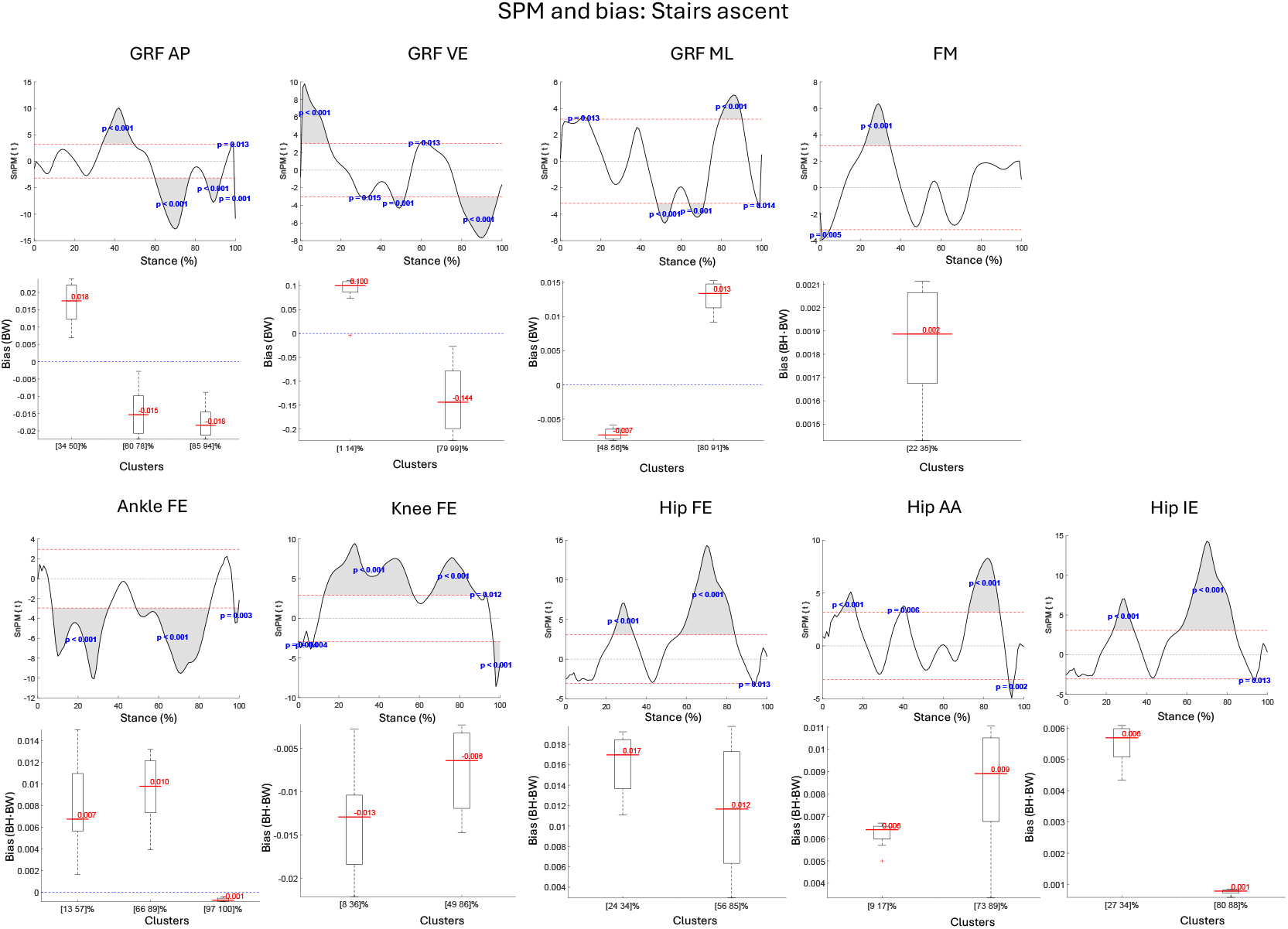
Results of SPM analysis between predictions and ground truth and related boxplots of biases for stance intervals with systematic differences during stairs ascent.

**Figure 10.**
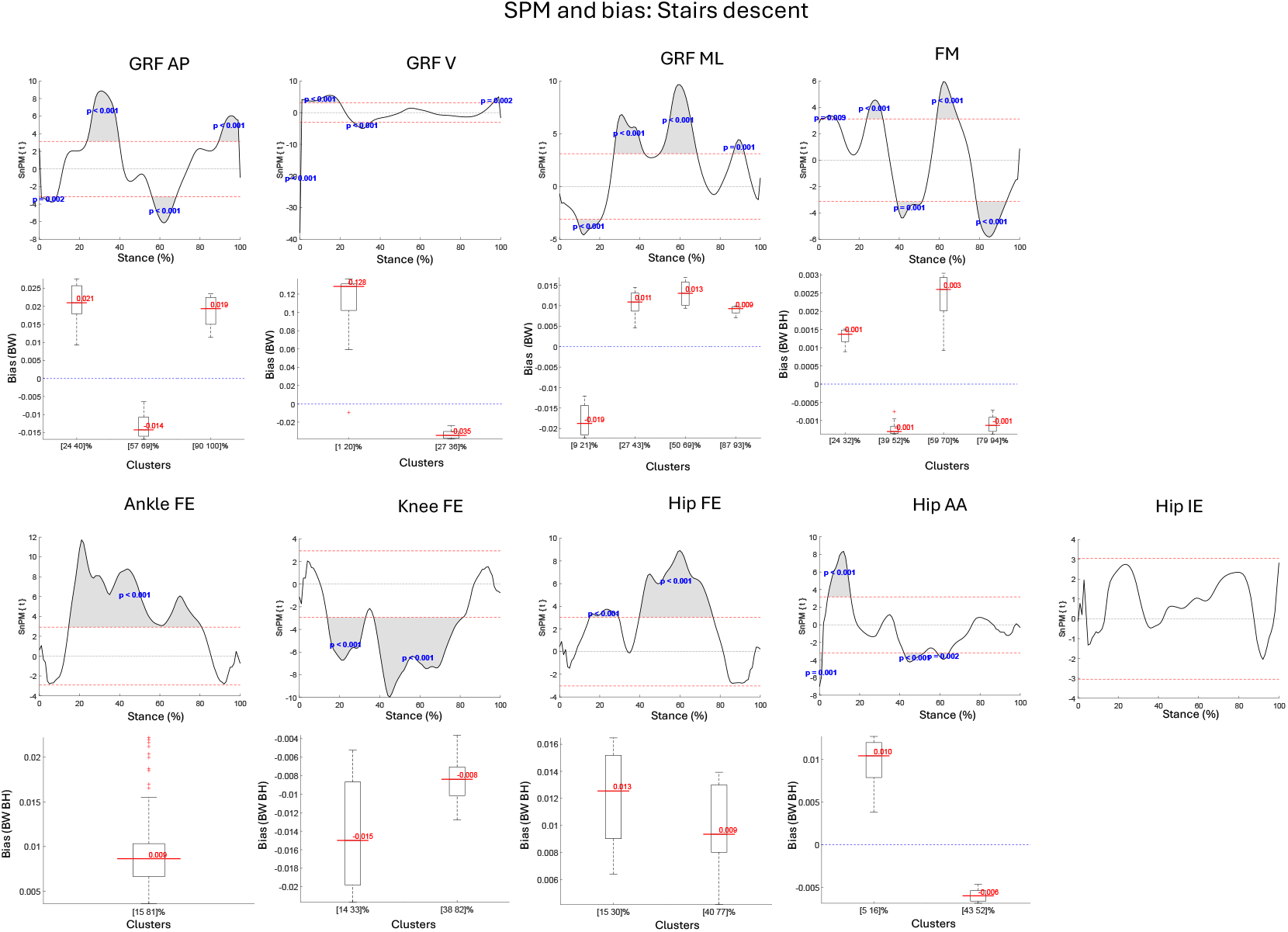
Results of SPM analysis between predictions and ground truth and related boxplots of biases for stance intervals with systematic differences during stairs descent.

For joint kinetics, the ankle flexion–extension torque was the most robust across tasks, with errors around 2% and correlation coefficient ρ>0.95. The knee flexion-extension moment was accurate in walking and stair ascent (nRMSE: 3.1– 8.3%, ρ up to 0.98) but less so in descent (nRMSE: 8.3 ± 4.2%, ρ = 0.71± 0.24). At the hip, abduction–adduction torque showed the highest accuracy across tasks, particularly in level walking, together with internal–external rotation moment (lowest nRMSE equal to 2.7%, highest ρ equal to 0.99), though larger errors were observed in stair ascent (nRMSE:10 ± 5.2%; ρ: 0.79 ± 0.11). In contrast, the hip flexion–extension torque was the least accurate, with moderate errors in walking (nRMSE: 7.1 ± 1.4% BH·BW; ρ: 0.78 ± 0.07) and the largest discrepancies in stair descent (nRMSE: 12.1 ± 5.0% BH·BW; ρ: 0.61 ± 0.19).

The bias quantification following SPM analysis revealed that the vertical GRF was slightly overestimated in early stance during walking (+0.11 BW, 1–8% of stance), strongly underestimated in late stance of stair ascent (–0.224 BW, 79–99%), and most overestimated in early stance of descent (+0.137 BW, 1–20%) (Table 2).

**Table 2.**
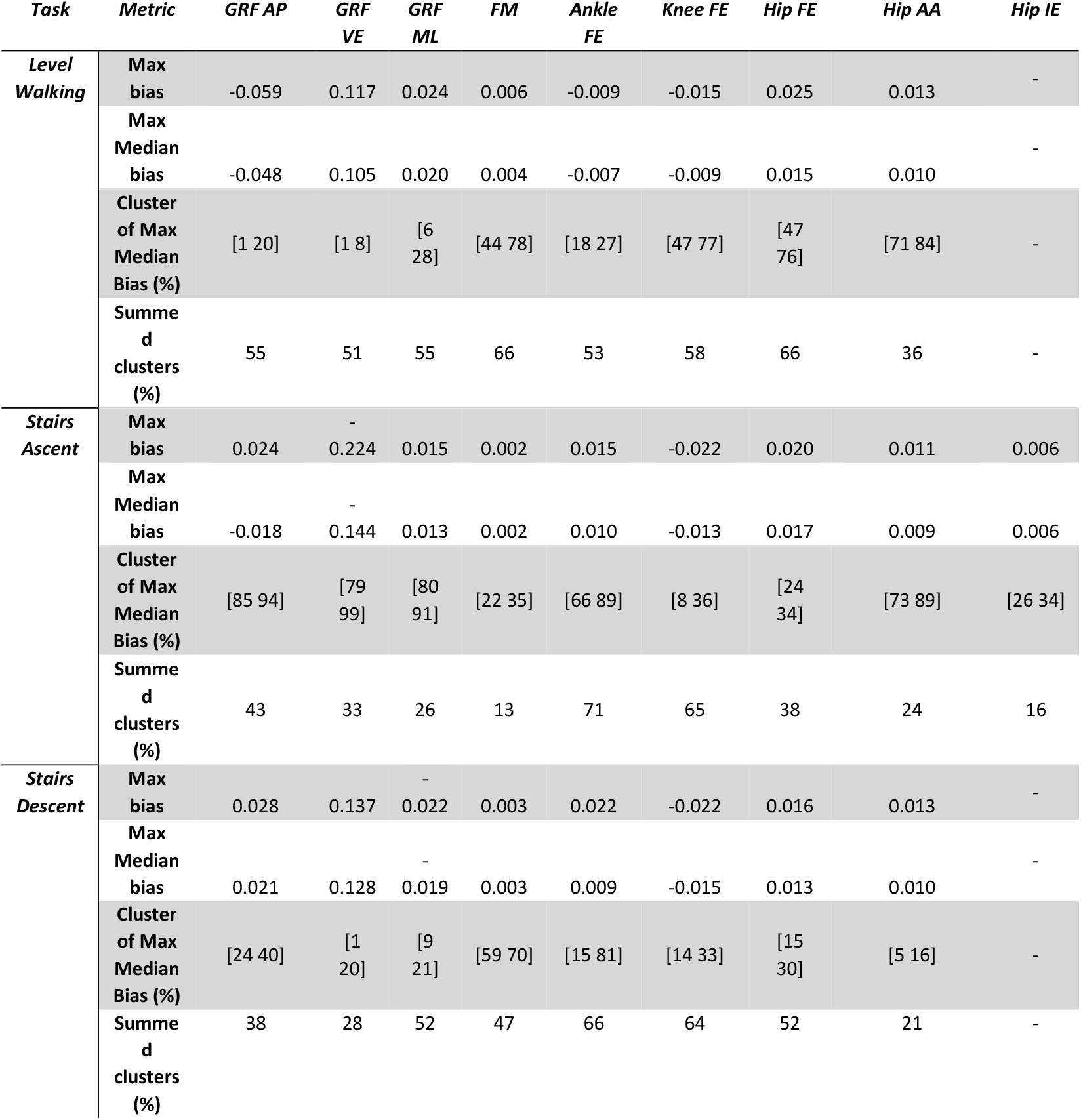
Results of SPM analysis showing the maximum bias over the entire stance phase (max bias), the maximum median bias (max median bias) within the related cluster (cluster of max median bias) and the total percentage of stance phase with systematically differences (summed clusters). Units of measurement are BW for GRF (ground reaction forces) and body height · body weight (BH·BW) for FM (free moment) and joint torques (AP=anterior-posterior, ML=medio-lateral, VE=vertical, FE=flexion-extension, AA=ab-adduction, IE=internal-external)

Regarding joint torques, ankle flexion–extension exhibited the most prolonged systematic errors over the entire task (i.e. summed clusters in Table 2) with overestimations of +0.015 BH·BW during stairs ascent, (for the 71% of stance phase) and +0.022 BH·BW during stairs descent (over the 66% of stance phase). Knee and hip flexion–extension moment biases achieved similar magnitudes with maximum values of -0.022 and 0.025 BH·BW, respectively in stairs ascent and level walking.

For CoP predictions, the largest errors were observed during stair descent (up to 140 mm) in the double-stance phase, where foot kinematics directly influence CoP estimation. Moreover, additional obstacles introduced by the experimental setup may have hindered the reconstruction of marker trajectories [39] which was also reflected in the increased maximum inverse kinematics error (up to 120 mm). Nonetheless, the overall predictions matched the experimental values satisfactorily, with median errors of 21, 23, and 21 mm for level walking, stair ascent, and stair descent, respectively. These values are comparable to the 24 mm of average RMSE reported by Podobnik et al.[40], adopting an approach of IMU-based statistical modelling.

## 4 Discussion and Conclusion

This study introduces and validates an optimization-based approach for predicting 3D GRF&Ms and CoP within the OpenSim environment [24], using only kinematic data and scaled MSK model. This methodology leverages the CMC algorithm to resolve double-support loading taking advantage of CoP estimates derived as FCP from ground-contact analysis. An open-source implementation of the proposed approach called OpenGRF is made available at https://simtk.org/projects/opengrf.

Validation of the proposed method against force plate data during level walking, stair ascent, and descent over 7 subjects (126 trials) demonstrated that OpenGRF provides robust and reliable GRF&M predictions across various locomotor tasks. Comparison with previous studies (Table 3) confirms this interpretation of the results. Remarkably low errors were achieved in level walking (nRMSE: 1.44% VE, 4.60% AP, 5.86% ML; ρ≤0.99), comparable to those reported for Fluit’s optimization-based framework, especially in the vertical component (3%). Fluit’s method yielded slightly larger errors in the AP direction with respect to the present study (15.8% vs 4.6%, in level walking), which may be related to the inclusion of muscle forces in the optimization process, potentially modifying the solution space and affecting in-plane GRF estimates. The subject- and trial-specific contact calibration implemented in OpenGRF appeared beneficial for enhancing consistency of the applied external loads location and may also contribute to the minor discrepancies observed between the two approaches. Both approaches had moderately accurate predictions of the free moment, which, as noted by Fluit et al.[1], may be due to the absence of a torsional knee DoF in hinge-type models limiting representation of transverse-plane motion.

**Table 3.**
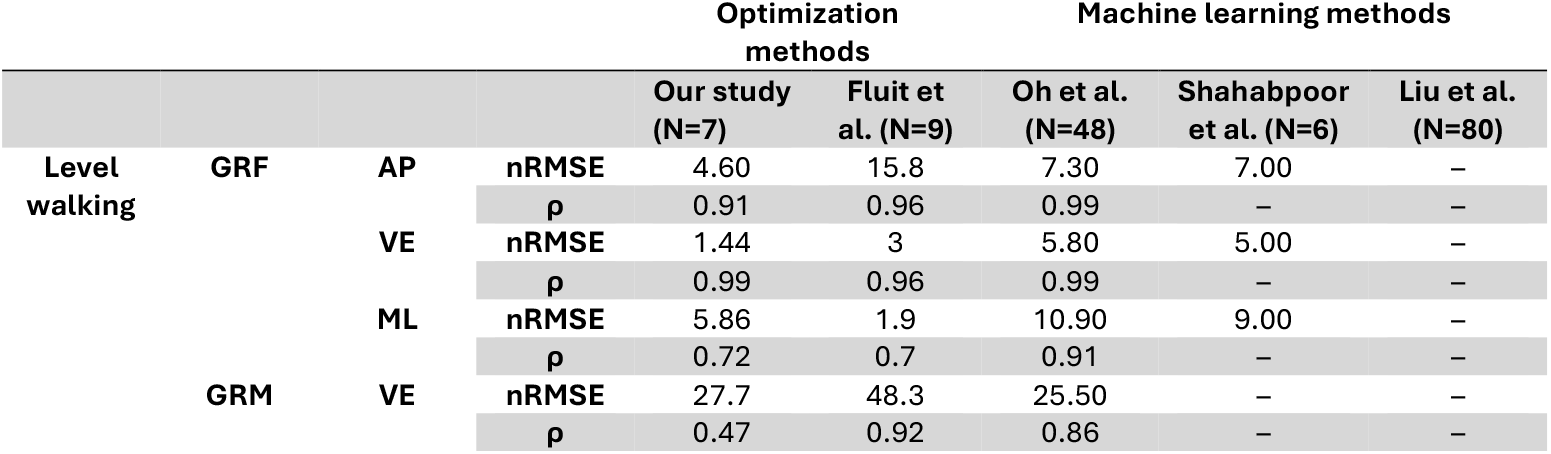
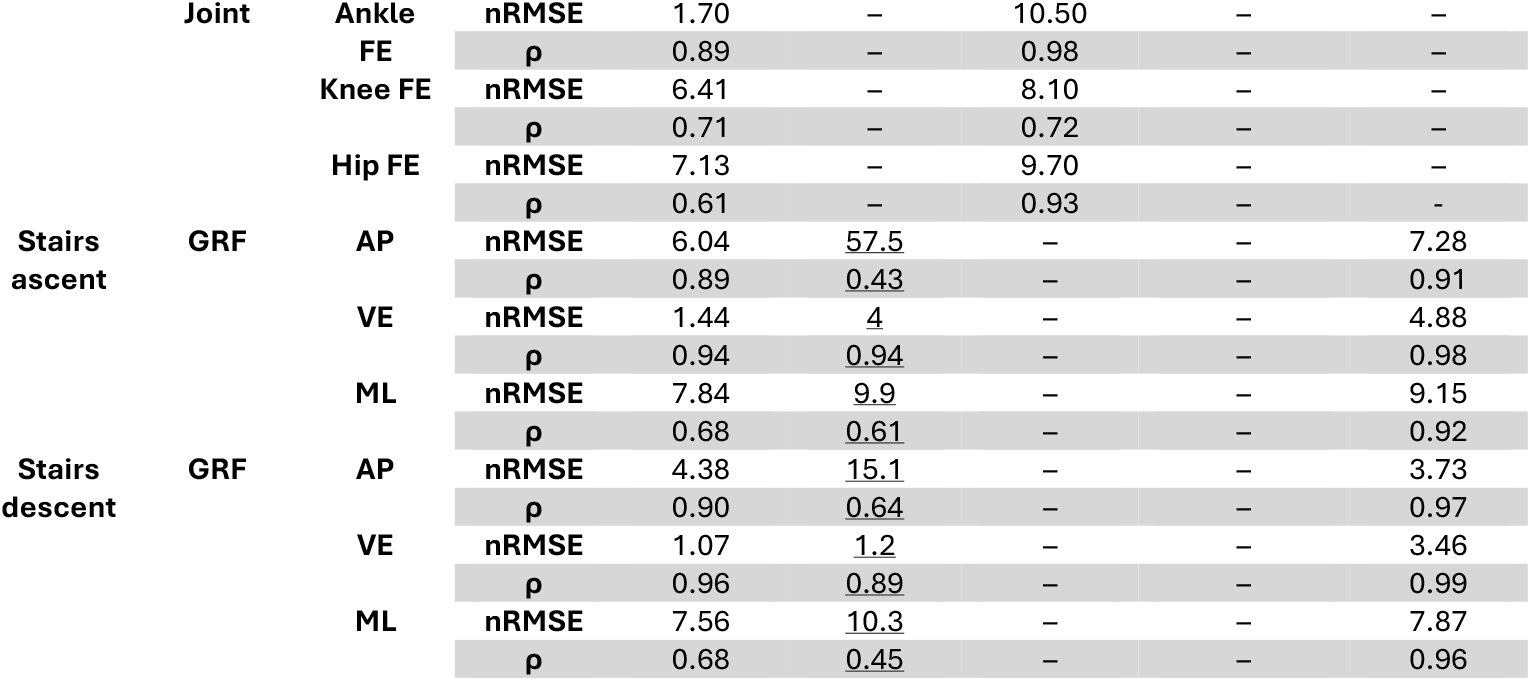
Summary of prediction accuracy (ρ and nRMSE) for various ground reaction forces and moments (GRF&M) methodologies. Undersigned values for Fluit et al. suggest that the values are mean errors.

Machine learning methods, such as Oh’s neural network [13] (including 48 subjects) and Liu’s BiLSTM model [19](comprising 80 patients), generally reported very high correlations (often ≥0.98), typically exceeding those of optimization-based approaches, reflecting their strong ability to reproduce waveform shapes once trained on sufficient data. However, these methods also showed larger nRMSE for certain components (e.g., Oh: 5.8% VE, 7.3% AP; Liu: up to 7–9% in stair ascent), likely suffering from the lack of a physics-based constraints, with performance closely linked to the size and characteristics of the training datasets. Similarly, Shahabpoor et al. [18] using IMUs (6 subjects) reported slightly higher errors with respect to the previous cited ones [13,19], and noted difficulties in capturing high-frequency GRF peaks, a limitation of IMU-derived kinematics compared to marker-based motion capture also emphasized by Tan et al. [41].

For CoP predictions, the largest errors were observed during stair descent (up to 140 mm) in the double-support phase, where foot kinematics directly influence CoP estimation. In stair negotiation tasks, an experimental limitation was the staircase occasionally occluding marker tracking [39] which was also reflected in the increased maximum inverse kinematics error (up to 120 mm). Nonetheless, the overall predictions matched the experimental values satisfactorily, with median errors of 21, 23, and 21 mm for level walking, stair ascent, and stair descent, respectively. These values are comparable to the 24 mm of average RMSE reported by Podobnik et al.[40], adopting an approach of IMU-based statistical modelling.

The SPM findings suggest that the proposed methodology presents a decreased accuracy when predicting abrupt vertical loading changes, particularly during transitional phases of stance in stair negotiation plausibly due to increased inertial forces that affect the load partitioning between the two limbs; accordingly, this behaviour causes misalignments, between predicted and reference waveforms, to which SPM is particularly sensitive [42], though not the nRMSE and ρ metrics.

It is also worth noting that GRF&M predictions in this study were performed with models excluding muscles, to reduce the dimensionality of the optimisation solution space. Incorporating muscle activations, as in the standard CMC algorithm, would have required defining appropriate relative weightings between the GRF&Ms prediction and muscle redundancy contributions to the objective function. Muscle recruitment can then be solved in a subsequent step, performing static optimization after the GRF&Ms have been estimated.

The presented methodology is subject to some limitations that should be considered when interpreting the results. First, modelling the foot as a single rigid segment introduces a simplification of the anatomical and functional complexity of the human foot, possibly affecting directly the FCP locations and accordingly the FM calculation during double stance phase. Second, the arbitrary contact probe’s locations on the feet, especially on heel and phalanges, straightly influences contact time intervals detection during the contact analysis; moreover, the calibration of contact probes is sensitive to both the scaled MSK model, and the input kinematic data. Specifically, the amount of cranial–caudal adjustment of the probes positions depends on the vertical gap between the foot and the contact plane at the foot-flat instant, since the algorithm shifts the probes to ensure consistent foot–ground penetration. Third, the GRF&M prediction accuracy strongly depends on the quality of the used MSK model, including segment lengths, masses and inertial properties [43]. Likewise, the quality of the input kinematics (e.g. limiting soft tissue artifacts [44] is critical for ensuring accurate predictions, and this could be a limiting aspect for combining this methodology with technologies characterised by larger uncertainties like markerless pose estimation [45]. Moreover, although the approach is in principle compatible with any full-body MSK model, the validation undertaken in this study was performed only with the FullBodyModel of Rajagopal et al. [31]; further validation is required when using different models, e.g. Gait2392 [46], Lai Arnold [47], in future work.

Future developments will focus on extending the tool’s validation to a broader range of motor tasks, including shod and inclined walking, dynamic movements [20,48] (e.g., jumps, squat jumps), and multi-support activities (e.g., Burpees [49], push-ups). In addition, extended validation with markerless motion capture approaches, such as OpenCap [50] or wearable IMUs [27], will be pursued to evaluate accuracy in on-field environments. Furthermore, assessing OpenGRF in a pathological population may provide valuable insights and serve as further validation of the tool’ robustness in clinical environment [22,51].

In conclusion, we presented a novel methodology, implemented in the OpenGRF tool, for predicting dynamics from kinematic data in OpenSim, achieving, and in some cases exceeding, state-of-the-art accuracy, particularly in the estimation of vertical ground reaction forces and ankle joint moments across a variety of daily living tasks.

## Abbreviations

GRFs: ground reaction forces
GRMs: ground Reaction Moment
GRF&Ms: ground reaction forces and moments
FM: free moment
CoP: centre of pressure
AP: anterior-posterior
VE: vertical
ML: medio-lateral
FE: flexion-extension
AA: ab-adduction
IE: internal-external
MSK: musculoskeletal
FCPnRMSE: normalized root mean square error
ρ: Pearson’s correlation coefficient
DoFs: degrees of freedom
FCP: foot contact point
SD: standard deviation
CMC: computed-muscle-control
SPM: statistical parametric mapping

## Authors contribution

Conceptualization: all authors; Data curation: ADP, LM; Formal analysis: ADP; Funding acquisition: LM, FDP; Investigation: ADP; Methodology: ADP; Project administration: all authors; Resources: FDP, LM; Software: ADP, LM Supervision: FDP, LM; Validation: all authors; Visualization: ADP; Writing - original draft: ADP, FDP; and Writing - review & editing: FDP, LM

## Funding acquisition

This work has been partly funded by the PNRR project “Tuscany Health Ecosystem” (THE) (Ecosistemi dell’Innovazione) - Spoke 6 - Precision Medicine & Personalized Healthcare (CUP I53C22000780001) under the NextGeneration EU programme and by a UNSW Scientia Fellowship to LM.

## Appendices

### A. Variables of the optimization problem

This section deepens the implementation of the ideal actuators into the optimization problem. The three sets of ideal actuators added to the model are:

1. Coordinate actuators: One per active DoF (excluding 6 pelvis DoFs), each with an optimal torque of 1000 Nm; these must balance the joint net moments required by the task.
2. Foot external actuators: one force acting at the FCP, with three directions (*d=1,3* means ground tangential components whilst *d=2* is the direction normal to the contact plane) and one torque (applied between the calcaneus and ground) with the same three directions; the vertical direction (d=2) represents a real “free moment”. The optimal external force is 10000 N to grant adequate support at feet; whilst the optimal value of GRM is 100 Nm.
3. Pelvis residual actuators: Six actuators between pelvis and ground (translational and rotational), each with optimal force/torque (1 N or 1 Nm) ease solution convergence.

### B. Variables constraints

This section summarizes the constant and time-varying constraints embedded in the optimization problem, expressed in section 2.2, for the above discussed variables:

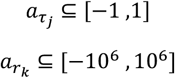

*If t* ∈ *CT1*:

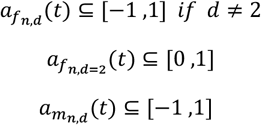

*If t* ∉ *CT1*:

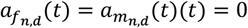

where the index *j* represents the model’s (internal) DoFs, excluding the pelvis DoFs indicated by *k*; *n* is the index of GRFs and GRMs for left and right foot while *d* refers to their components (d=2 means the direction normal to the contact level).

### C. Theoretical background for CoP calculation

For a system of particles or rigid bodies having centre of mass in CoM, Newtonian equations of dynamics state that:

1. The sum of the external forces acting on a system is equal to the product of its total mass and the acceleration of G:

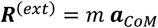

And
2. The total external force moment with respect to CoM, ***M***_*CoM*_^(*ext*)^ is equal to the time derivative of angular momentum with respect to CoM (***L***_*CoM*_):

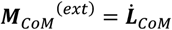

When we consider the human body as a system interacting with the environment through the feet, the external forces and moments are given by the GRFs and GRMs, thus the previous equations become:

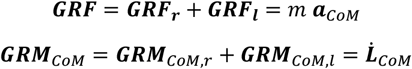

where the subscript l and r are used to distinguish the actions from the left and right foot.

In a musculoskeletal analysis, the right-end side of both equations depend on the kinematics and inertial properties of the system, which are usually estimated by scaling a model and solving the inverse kinematic problem. Therefore, dynamics equations can be used to calculate the resultant ground reaction forces and moments (***GRF*** and ***GRM***_*CoM*_), for example when force plate data are not available. It comes evident that the critical point is to distinguish the contribution of each foot when both are in contact with the ground, as the problem becomes indeterminate. Additional criteria/approaches should be introduced to overcome this issue.

Another point is worth being clarified: the point of application of the **GRF**s. It is common practice to consider GRF applied at the CoP of each foot, defined as the point of ground with respect to the GRMs have only a vertical component (the free moment, *FM*). Once ***GRM***_*CoM*_ is known, the CoP location, its two coordinates in the ground plane and FM can be calculated from:

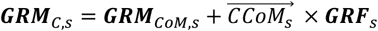

with s=l,r and C=CoP. Thus, to correctly place the feet CoPs, it is necessary to have the separate contribution of GRM for each foot. The above equation can be used also to find the moment with respect to other convenient point.

### D. CoP weight function

In this section we have reported the weight function (w) to ensure continuity in the CoP trajectories across single and double stance phases (Figure 3). It is a piecewise defined function applied to the CoP trajectories during the stance phase (Eq. 7):

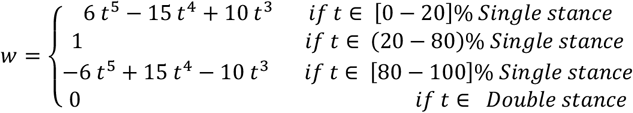

Where *t* is the time.

### E. Tracking errors

The values in Table 4 represent tracking errors (°) for the model lower-body coordinates between experimental input kinematics and the simulated kinematics obtained through CMC. Global errors among subjects are reported as mean (SD) across level gait, stair ascent, and stair descent. Overall, errors were small, generally below 1° for the pelvis and lumbar segments, while hip and knee angles showed mean errors between 0.2° and 1.3°. The largest deviations were observed at the ankle (up to 2.1°) and subtalar joints (up to 2.3°), slightly exceeding the recommended accuracy in some case (< 2°), but still generally considered acceptable for tracking [36].

**Table 4.** Tracking errors in degrees (°) for the model lower-body coordinates between input kinematics and the one in output from CMC simulations, averaged (mean (SD)) across all the subjects and motor tasks.

| Pelvis |  | Hip & Knee |  | Ankle & Subtalar |  | Lumbar |  |
| --- | --- | --- | --- | --- | --- | --- | --- |
| Variable | Mean (SD) | Variable | Mean (SD) | Variable | Mean (SD) | Variable | Mean (SD) |
| Pelvis tilt | 0.29 (0.16) | Hip flexion | 0.57 (0.29) | Ankle angle | 1.49 (0.62) | Lumbar extension | 0.36 (0.2) |
| Pelvis list | 0.33 (0.16) | Hip adduction | 0.45 (0.19) | Subtalar angle | 1.340 (1.03) | Lumbar bending | 0.39 (0.2) |
| Pelvis rotation | 0.24 (0.11) | Hip rotation | 0.85 (0.61) |  |  | Lumbar rotation | 0.29 (0.13) |
| Pelvis AP | <0.01 | Knee angle | 0.9 (0.45) |  |  |  |  |
| Pelvis SI | <0.01 |  |  |  |  |  |  |
| Pelvis ML | <0.01 |  |  |  |  |  |  |

### F. Residuals at pelvis

The values reported in Table 5 correspond to the mean and SD of the RMS residuals obtained across all level walking, stair ascent, and stair descent trials. Although they do not physically represent discrepancies with respect to experimental data (as typically occurs when introducing experimental GRF&Ms), these residuals at pelvis remained widely inside the tolerance limits (10 N for forces and 30 Nm for moments) reported in the CMC online documentation [36]

**Table 5.** RMS pelvis residuals in terms of mean (SD) between all the gait, stair ascent, and stair descent trials

| RMS Pelvis Residual | Mean (SD) |
| --- | --- |
| Tilt (Nm) | 1.38 (3.32) |
| List (Nm) | 0.15 (0.64) |
| Rotation (Nm) | 0.03 (0.09) |
| Anterior-Posterior (N) | 1.26 (3.03) |
| Superior-Inferior (N) | 0.44 (1.16) |
| Medio-Lateral (N) | 0.14 (0.64) |

**Table 6.** Pearson’s correlation coefficient (ρ) in terms of mean (SD) between GRF&M predicted and ground truth for all the subjects and tasks averaged across trials

| Task | Subj | GRF AP | GRF VE | GRF ML | FM | Ankle FE | Knee FE | Hip FE | Hip AA | Hip IE |
| --- | --- | --- | --- | --- | --- | --- | --- | --- | --- | --- |
| Level Walking | 1 | 0.969 | 0.983 | 0.749 | 0.746 | 0.979 | 0.384 | 0.768 | 0.867 | 0.845 |
|  |  | (0.011) | (0.006) | (0.041) | (0.082) | (0.010) | (0.223) | (0.026) | (0.047) | (0.083) |
|  | 2 | 0.942 | 0.939 | 0.857 | 0.780 | 0.983 | 0.848 | 0.695 | 0.880 | 0.916 |
|  |  | (0.008) | (0.021) | (0.047) | (0.070) | (0.008) | (0.062) | (0.012) | (0.026) | (0.017) |
|  | 3 | 0.868 | 0.919 | 0.817 | 0.708 | 0.970 | 0.667 | 0.712 | 0.901 | 0.687 |
|  |  | (0.024) | (0.020) | (0.045) | (0.122) | (0.012) | (0.083) | (0.081) | (0.018) | (0.105) |
|  | 4 | 0.951 | 0.912 | 0.659 | 0.916 | 0.990 | 0.875 | 0.838 | 0.811 | 0.773 |
|  |  | (0.014) | (0.014) | (0.052) | (0.026) | (0.006) | (0.045) | (0.009) | (0.058) | (0.024) |
|  | 5 | 0.905 | 0.940 | 0.797 | 0.697 | 0.982 | 0.771 | 0.851 | 0.913 | 0.917 |
|  |  | (0.018) | (0.014) | (0.051) | (0.178) | (0.002) | (0.054) | (0.021) | (0.023) | (0.016) |
|  | 6 | 0.775 | 0.908 | 0.533 | 0.492 | 0.978 | 0.623 | 0.822 | 0.834 | 0.900 |
|  |  | (0.024) | (0.011) | (0.133) | (0.096) | (0.005) | (0.139) | (0.025) | (0.035) | (0.025) |
|  | 7 | 0.980 | 0.953 | 0.680 | 0.144 | 0.974 | 0.900 | 0.793 | 0.873 | 0.911 |
|  |  | (0.007) | (0.012) | (0.156) | (0.192) | (0.009) | (0.011) | (0.030) | (0.034) | (0.032) |
| Stairs ascent | 1 | 0.764 | 0.917 | 0.498 | 0.468 | 0.951 | 0.954 | 0.618 | 0.828 | 0.815 |
|  |  | (0.057) | (0.020) | (0.148) | (0.082) | (0.022) | (0.019) | (0.105) | (0.055) | (0.065) |
|  | 2 | 0.751 | 0.952 | 0.765 | 0.595 | 0.962 | 0.968 | 0.636 | 0.744 | 0.854 |
|  |  | (0.116) | (0.029) | (0.092) | (0.087) | (0.015) | (0.013) | (0.144) | (0.038) | (0.028) |
|  | 3 | 0.954 | 0.989 | 0.851 | 0.185 | 0.954 | 0.964 | 0.583 | 0.903 | 0.743 |
|  |  | (0.011) | (0.005) | (0.067) | (0.141) | (0.025) | (0.016) | (0.235) | (0.054) | (0.129) |
|  | 4 | 0.745 | 0.940 | 0.607 | 0.264 | 0.940 | 0.944 | 0.446 | 0.803 | 0.706 |
|  |  | (0.171) | (0.018) | (0.275) | (0.116) | (0.026) | (0.011) | (0.249) | (0.095) | (0.076) |
|  | 5 | 0.827 | 0.922 | 0.589 | 0.392 | 0.936 | 0.966 | 0.632 | 0.856 | 0.726 |
|  |  | (0.032) | (0.061) | (0.193) | (0.123) | (0.047) | (0.009) | (0.154) | (0.020) | (0.116) |
|  | 6 | 0.810 | 0.966 | 0.787 | 0.714 | 0.959 | 0.943 | 0.642 | 0.881 | 0.928 |
|  |  | (0.038) | (0.016) | (0.046) | (0.062) | (0.013) | (0.014) | (0.079) | (0.046) | (0.020) |
|  | 7 | 0.814 | 0.861 | 0.661 | 0.681 | 0.929 | 0.898 | 0.252 | 0.777 | 0.773 |
|  |  | (0.029) | (0.053) | (0.148) | (0.099) | (0.031) | (0.036) | (0.103) | (0.074) | (0.044) |
| Stairs descent | 1 | 0.704 | 0.919 | 0.466 | 0.407 | 0.788 | 0.416 | 0.444 | 0.615 | 0.645 |
|  |  | (0.261) | (0.083) | (0.123) | (0.228) | (0.161) | (0.174) | (0.267) | (0.211) | (0.155) |
|  | 2 | 0.926 | 0.972 | 0.754 | 0.417 | 0.880 | 0.976 | 0.854 | 0.841 | 0.928 |
|  |  | (0.012) | (0.015) | (0.066) | (0.104) | (0.048) | (0.006) | (0.048) | (0.028) | (0.034) |
|  | 3 | 0.931 | 0.906 | 0.824 | 0.342 | 0.847 | 0.359 | 0.508 | 0.906 | 0.848 |
|  |  | (0.027) | (0.047) | (0.050) | (0.346) | (0.061) | (0.132) | (0.059) | (0.032) | (0.043) |
|  | 4 | 0.939 | 0.972 | 0.837 | 0.288 | 0.958 | 0.810 | 0.618 | 0.954 | 0.848 |
|  |  | (0.018) | (0.018) | (0.037) | (0.146) | (0.022) | (0.086) | (0.131) | (0.011) | (0.126) |
|  | 5 | 0.933 | 0.990 | 0.488 | 0.382 | 0.900 | 0.714 | 0.571 | 0.940 | 0.818 |
|  |  | (0.014) | (0.002) | (0.285) | (0.189) | (0.044) | (0.080) | (0.133) | (0.040) | (0.077) |
|  | 6 | 0.924 | 0.991 | 0.741 | 0.373 | 0.891 | 0.773 | 0.544 | 0.901 | 0.837 |
|  |  | (0.033) | (0.006) | (0.087) | (0.127) | (0.026) | (0.127) | (0.091) | (0.016) | (0.101) |
|  | 7 | 0.943 | 0.988 | 0.705 | 0.553 | 0.942 | 0.907 | 0.693 | 0.937 | 0.885 |
|  |  | (0.039) | (0.006) | (0.190) | (0.178) | (0.029) | (0.019) | (0.157) | (0.026) | (0.044) |

**Table 7.**
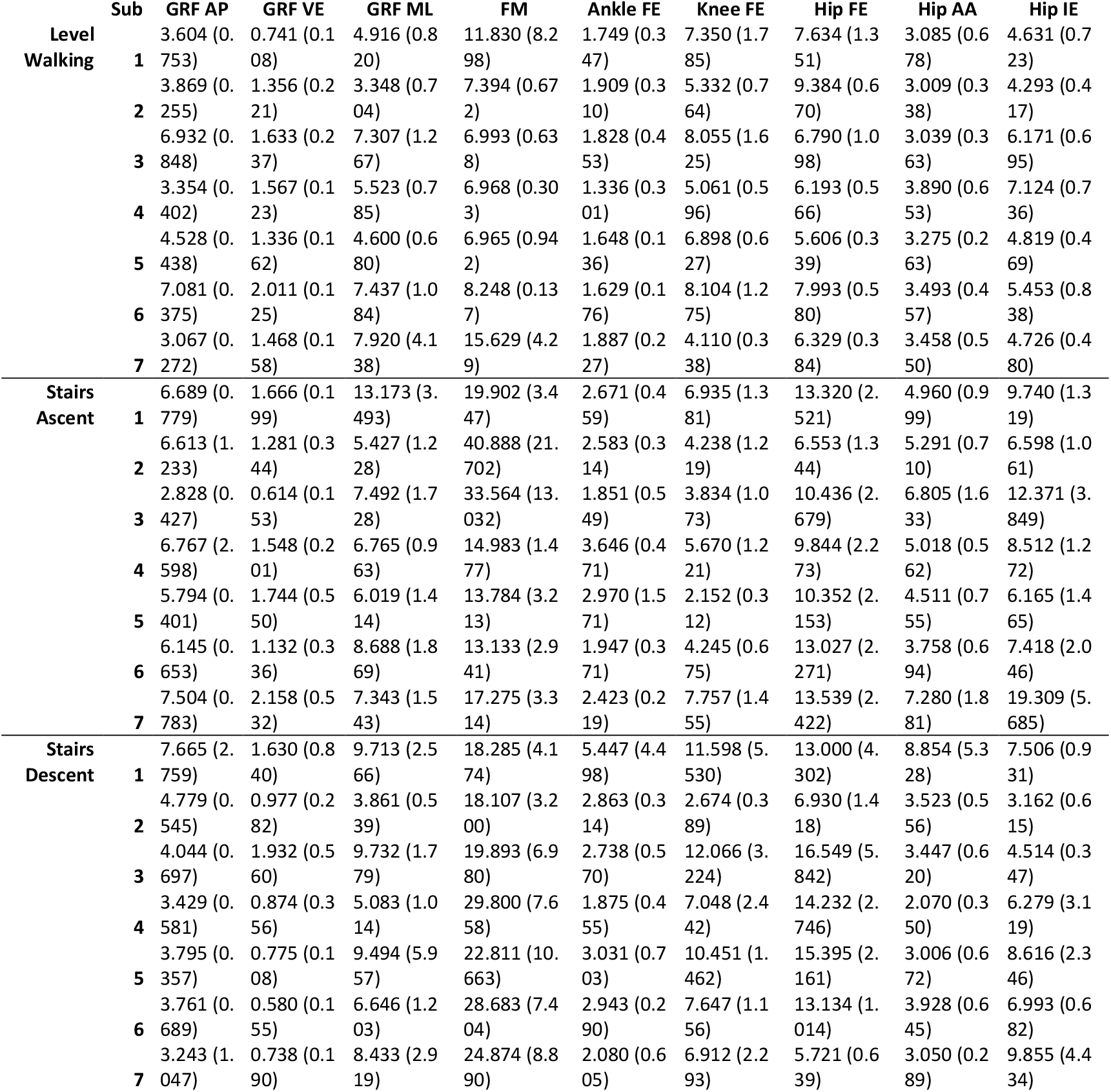
mean (SD) normalized root mean square error (nRMSE) between GRF&M predicted and ground truth for all subjects and tasks averaged across trials

### G. Supplementary materials

This section shows an example of the results obtained for SPM and bias analysis (shown in Results, Table 4). Due to the large number of resulting plots, only the one related to vertical GRF during level walking is presented.

